# Fast assessment of human receptor-binding capability of 2019 novel coronavirus (2019-nCoV)

**DOI:** 10.1101/2020.02.01.930537

**Authors:** Qiang Huang, Andreas Herrmann

**Author notes:** To whom correspondence should be addressed. (Q.H.); (A.H.).

## Abstract

The outbreaks of 2002/2003 SARS, 2012/2015 MERS and 2019/2020 Wuhan respiratory syndrome clearly indicate that genome evolution of an animal coronavirus (CoV) may enable it to acquire human transmission ability, and thereby to cause serious threats to global public health. It is widely accepted that CoV human transmission is driven by the interactions of its spike protein (S-protein) with human receptor on host cell surface; so, quantitative evaluation of these interactions may be used to assess the human transmission capability of CoVs. However, quantitative methods directly using viral genome data are still lacking. Here, we perform large-scale protein-protein docking to quantify the interactions of 2019-nCoV S-protein receptor-binding domain (S-RBD) with human receptor ACE2, based on experimental SARS-CoV S-RBD-ACE2 complex structure. By sampling a large number of thermodynamically probable binding conformations with Monte Carlo algorithm, this approach successfully identified the experimental complex structure as the lowest-energy receptor-binding conformations, and hence established an experiment-based strength reference for evaluating the receptor-binding affinity of 2019-nCoV via comparison with SARS-CoV. Our results show that this binding affinity is about 73% of that of SARS-CoV, supporting that 2019-nCoV may cause human transmission similar to that of SARS-CoV. Thus, this study presents a method for rapidly assessing the human transmission capability of a newly emerged CoV and its mutant strains, and demonstrates that post-genome analysis of protein-protein interactions may provide early scientific guidance for viral prevention and control.

## Introduction

Coronaviruses (CoV) are positive-stranded enveloped RNA viruses belonging to the order *Nidovirales*, and are classified into four genera: α, β, δ and γ. Two of the β-CoVs, the server acute respiratory syndrome CoV (SARS-CoV) and the Middle East respiratory syndrome CoV (MERS-CoV), have caused serious epidemics. In November 2002, SARS emerged in southern China and then was spreading across the world in 2003, resulting in ~8,000 cases with a fatality rate of about 10%(WHO, 2004). Ten years later, MERS emerged in Saudi Arabia and spread to different countries with a fatality rate of about 35%(WHO, 2016). Both viruses are of zoonotic nature: SARS-CoV originates from bats/civet cats(Ge et al., 2013), and the major reservoir for MERS-CoV is dromedary camels(Reusken et al., 2016). Nevertheless, both viruses can be transmitted from human to human(de Wit et al., 2016). In contrast, a few other human CoVs of zoonotic origin only cause mild infections of the respiratory tract, such as HCoV-OC43 and HCoV-HKU1(Gaunt et al., 2010).

In December 2019, a novel CoV (2019-nCoV) emerged from Wuhan, China, and spread rapidly to other areas(Holshue et al., 2020; Li et al., 2020; Wu et al., 2020; Zhou et al., 2020). Although its original host remains unknown, all available data point again to a wild animal source. And, it is evident that human-to-human transmission is causing the rapid spreading of this newly emerged virus. Viral strains from infected persons of the area have been sequenced; but only little genetic variation was found, implying that they are descended from a common ancestor(Rambaut, 2020). Full-length genome analysis of 2019-nCoV patient samples revealed 79.5% and 96% sequence identities to SARS-CoV and a bat SARSr-Cov (severe acute respiratory syndrome-related CoV), respectively; however, sequences of the seven conserved viral replicase domains in ORF1ab show 94.6% identity between 2019-nCoV and SARS-CoV(Zhou et al., 2020).

It is widely accepted that the CoV human transmission is driven by the interactions of its trimeric transmembrane spike glycoprotein (S-protein) with the human receptors on the host cell surface. The S-protein consists of two subunits (i.e., S1 and S2) and is essential for the early steps of host cell infection: S-protein of a virion binds to the host cell receptors via its S1 receptor-binding domain (S-RBD) and then fuses the viral membrane with cellular membranes. The host cell receptors differ among human CoVs: SARS-CoV binds to exopeptidase angiotensin-converting enzyme 2 (ACE2)(Li et al., 2005; Li et al., 2003), and MERS-CoV to dipeptidyl peptidase 4(DPP4)(Raj et al., 2013); other CoVs such as HCoV-HKU1 use O-acetylated sialic acid as a receptor(Huang et al., 2015).

Sequence alignment of S1-proteins has revealed a sequence identity of about 75% between that of 2019-nCoV and those of several other CoVs including SARS-CoV(Zhou et al., 2020). Comparing with that of SARS-CoV, the S-RBD sequence of 2019-nCoV S-protein is very conserved, implying that ACE2 is the specific receptor for this virus to bind to the human host cells(Lu et al., 2020; Xu et al., 2020). Indeed, recent experimental evidences also supported that ACE2 is the human receptor of 2019-nCoV(Zhou et al., 2020). Consequently, it becomes very interesting to characterize the human receptor-binding capability of 2019-nCoV by analyzing the S-protein interactions with ACE2, in order to assess its potential for the human-to-human transmission.

Here, we employ protein-protein docking approach to quantify the interactions of 2019-nCoV S-RBD with ACE2, based on the experimentally determined structure of SARS-CoV S-RBD complexed with ACE2(Li et al., 2005) and on the predicted structure of 2019-nCoV S-RBD according to gene sequence. Computational protein-protein docking with accurate, physics-based energy functions is able to reveal the native-like, low-energy protein-protein complex from the unbound structures of two individual, interacting protein components(Chaudhury et al., 2011), such as antibody-antigen structures(Weitzner et al., 2017), and thereby may calculate the binding free energy of the investigated protein-protein interactions. To this end, large-scale local protein-protein docking was first carried out to sample a large number of thermodynamically probable conformations for the S-RBD binding to ACE2. Next, the binding free energy of S-RBD to ACE2 was calculated according to the thermodynamic average over the sampled, low-energy binding conformations. Our results show that the human receptor-binding affinity of 2019-nCoV is about 73% of that of SARS-CoV, implying that 2019-nCoV is able to bind to ACE2, and thereby infects the human host cells like SARS-CoV.

## Results and Discussion

### Binding interactions of SARS-CoV S-RBD with human ACE2

To quantify the binding interactions of CoV S-protein with ACE2, we employed the well-established method of local protein-protein docking implemented in Rosetta software suite(Das and Baker, 2008). Using this approach, we sampled a large number of low-energy conformations of S-RBD in complex with ACE2 by preforming multiple, independent docking runs (i.e., simulations), and calculated their binding energies with a physically realistic, all-atom energy function for macromolecular modeling and design(Alford et al., 2017). According to statistical thermodynamics, the sampled conformations are considered to be accessible in the real-world process of S-protein binding to ACE2, depending on their thermodynamic probabilities determined by the binding energies.

To validate the docking approach, we first carried out local protein-protein docking to investigate the binding interactions of SARS-CoV S-RBD with ACE2, in order to test whether this approach could identify the lowest-energy conformations of S-RBD complexed with ACE2 identical or close to the experimental structure(Li et al., 2005). The latter is considered to be the most probable, native-like binding conformation in reality. As described in Methods, similar to the studies of Chaudhury et al. (2011), the co-crystal structure of SARS-CoV S-RBD in complex with ACE2 (PDB 2AJF) was used as the starting complex for the local protein-protein docking. Then, more than 3,000 independent docking runs were conducted to sample the low-energy binding conformations by optimizing rigid-body orientation and side-chain conformations, according to the docking protocols in Methods. The binding free-energy scores of the obtained complex models are shown in the scatter plot in Fig. 1A, against their RMSD (root mean square deviation) values of interface residues with respect to those in the starting co-crystal structure.

**Figure 1.**
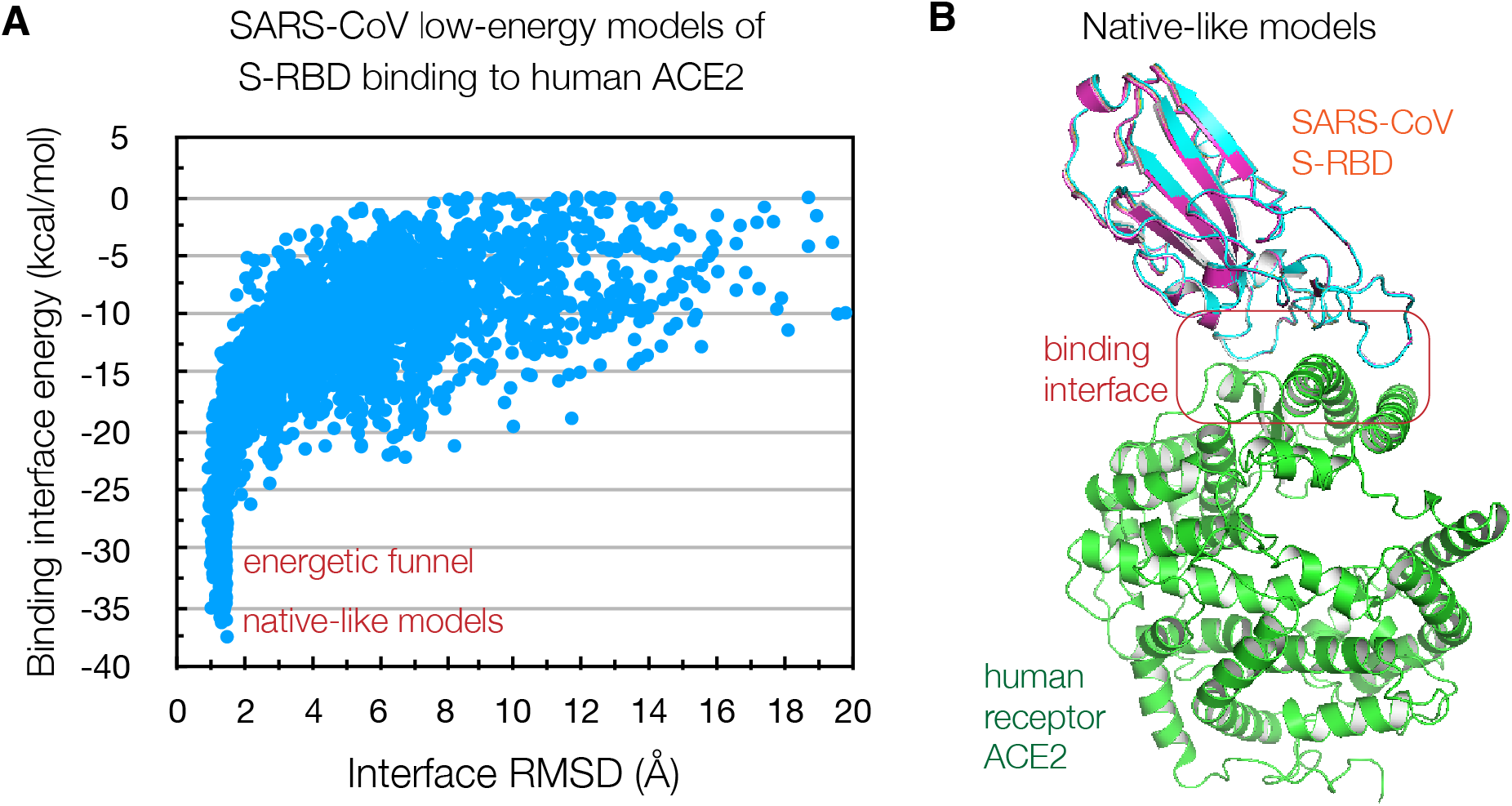
Binding interactions between SARS-CoV S-RDB and the human receptor ACE2. (A) Scatter plot of Rosetta binding free-energy scores versus RMSDs of interface residues of about 3,000 thermodynamically probable, low-energy binding models, where the RMSD reference is the experimental complex structure (PDB 2AJF). (B) The five top-scoring binding models. They are almost identical to the experimental structure with RMSDs < 1.5 Å.

As seen, a deep energetic funnel exhibits in the plot, and therefore clearly indicates that the docking simulations are very successful, because the presence of such a funnel is the most robust measure of successful docking simulations(Chaudhury et al., 2011). Due to their locations at the funnel bottom, the top-scoring models have better binding free-energy scores than other low-energy models, and therefore are considered to be the most probable, native-like binding conformations in thermodynamics. Significantly, the number of the native-like models among the top five scores (*N*_5_) is 5; also, their interface RMSDs are within 1.5 Å. Therefore, the lowest-energy conformations identified by the docking approach are almost identical to the experimental structure (PDB 2AJF). This provides a very solid basis for using this approach to investigate the S-RBD-ACE2 interactions of 2019-nCoV, in which no experimental complex structure is available because of time limitation, but only the first-determined genome data that can be used to predict the 2019-nCoV S-RBD structure.

The successful identification of the experimental complex structure as the lowest-energy binding conformations also validates the accuracy of the used Rosetta all-atom energy function(Alford et al., 2017) for quantifying the interactions of S-RBD with ACE2. As we will show, we could use the thermodynamic average of the binding energy scores of these conformations as an experiment-based energy reference to characterize the interactions of the 2019-nCoV S-RBD binding to ACE2, and then to evaluate the relative strength with respect to that of SARS-CoV. Such a comparison assessment avoids using the calculated, absolute free-energy scores to correlate with the binding strength, which might systematically differ from the actual measurement values because of the intrinsic, theoretical approximations of the used Rosetta all-atom energy function(Alford et al., 2017). We want to point out that all the docking simulations for SARS-CoV and 2019-nCoV use the same protocols and a consistent all-atom energy function. So, even the calculated energy scores contain certain errors caused by theoretical approximations, determination of the relative binding strength by comparing the calculated energy scores of two viruses should be fairly accurate.

### High-resolution prediction of 2019-nCoV S-RBD structure

Because the 2019-nCoVgenome has just been sequenced for a very short period of time, an experimental 3D structure for performing protein-protein docking of 2019-nCoV S-RBD to ACE2 is not yet available. Alternatively, computational methods may be used to predict the 3D structure(Schwede et al., 2009). Fortunately, the 2019-nCoV genome has shown that the amino-acid sequence of the S-protein is about 75% identical to that of the SARS-CoV S-protein (Fig. 2A). Therefore, with the SARS-CoV S-RBD structure as a template, it is possible to use high-resolution homology modeling method to accurately predict the 3D model(Qian et al., 2007).

**Figure 2.**
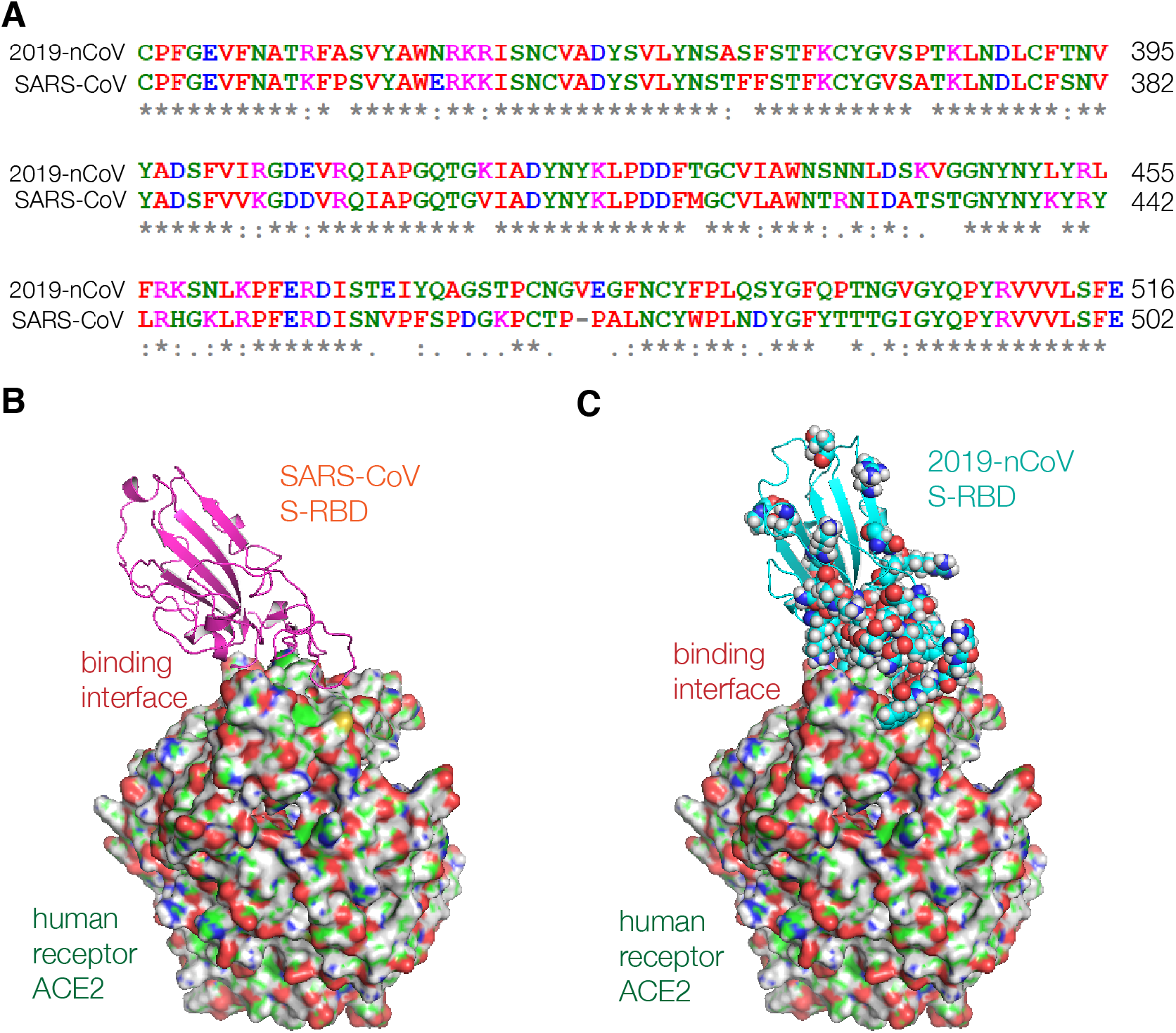
Homology modeling of 2019-nCoV S-RBD structure. (A) Amino-acid sequence alignment of 2019-CoV and SARS-CoV S-RBDs with an identity of 72.8%. (B) Experimental SARS-CoV S-RBD-ACE2 complex (PDB 2AJF). ACE2 is shown as surface model. (C) Starting complex of 2019-nCoV S-RBD-ACE2 for docking. For comparison, about 50 residues that are different from those of SARS-CoV S-RBD are shown as sphere models (see also Fig. 2A).

To the end, we first generated an initial model for the 2019-nCoV S-RBD via SWISS-MDOEL server at https://swissmodel.expasy.org, with the experimental S-RBD structure of SARS-CoV as the template (PDB 2AJF). Then, we performed all-atom refinements to optimize this initial model with Rosetta protocols (see Methods). The top-1 scoring model was used as the 3D structure of the 2019-nCoV S-RBD for the protein-protein docking with the experimental structure of ACE2 from PDB 2AJF. By superimposing this model onto the SARS-CoV S-RBD structure (Fig. 2B), we obtained the starting configuration of S-RBD complexed with ACE2 for the 2019-nCoV docking (Fig. 2C). From this complex, we found that most of the residues that are different from those of SARS-CoV S-RBD are close to or located in the binding interface. No doubt, such a structural distribution of the residues could lead to a different binding affinity from that of SARS-CoV S-RBD binding to ACE2.

### Binding interactions of 2019-nCoV S-RBD with human ACE2

With the complex model in Fig. 2C, more than 3,000 independent docking runs were performed to identify thermodynamically probable, low-energy binding conformations. The results are illustrated in Fig. 3. As shown in the scatter plot in Fig. 3A, the docking simulations also achieved an energetic funnel, in which four models among the top-5 scores have interface RMSDs within 0.7 Å. This demonstrates that the docking has identified the native-like interacting conformations for the 2019-nCoV S-RBD binding to ACE2. We consider these lowest-energy conformations at the energetic funnel bottom as those being engaged in the receptor-binding function.

**Figure 3.**
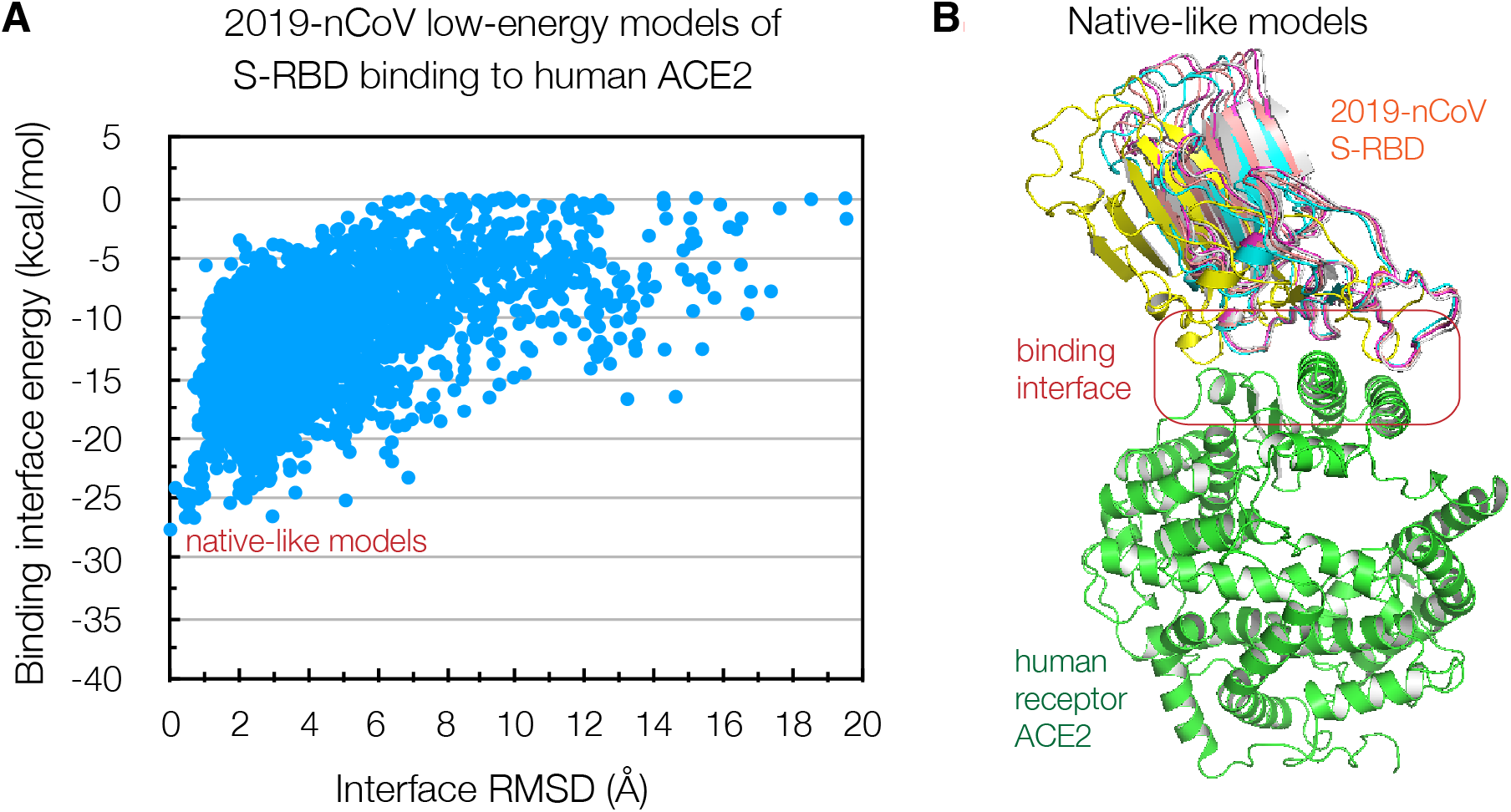
Binding interactions between 2019-nCoV S-RBD and the human receptor ACE2. (A) Scatter plot of Rosetta binding free-energy scores versus RMSDs of interface residues of about 3,000 thermodynamically probable, low-energy binding models, where the RMSD reference is the lowest-energy conformation. (B) The five top-scoring binding models.

When comparing with the SARS-CoV results in Fig. 1A, we found that there is no binding conformation with a binding energy score less than that of the lowest-energy conformation of SARS-CoV derived from experimentally determined structure (i.e., −37.4 kcal·mol^-1^ in Fig. 1A). To be sure, we also carried out high-resolution all-atom refinements to further optimize the lowest-energy binding conformation by 200 independent runs (see Methods). As indicated the scatter plot in Fig. 4A, the lowest-energy value of the 200 refined models remains about −28.4 kcal·mol^-1^ (Fig. 4B). Considering that the protein-protein docking has already sampled a large number of thermodynamically probable, low-energy binding conformations, these results indicate that the existence of an ACE2-binding conformation for 2019-nCoV S-RBD with an energy value < −37.4 kcalomol^-1^ is very limited. So, the binding strength of 2019-nCoV S-RBD to ACE2 is unlikely greater than that of SARS-CoV S-RBD.

**Figure 4.**
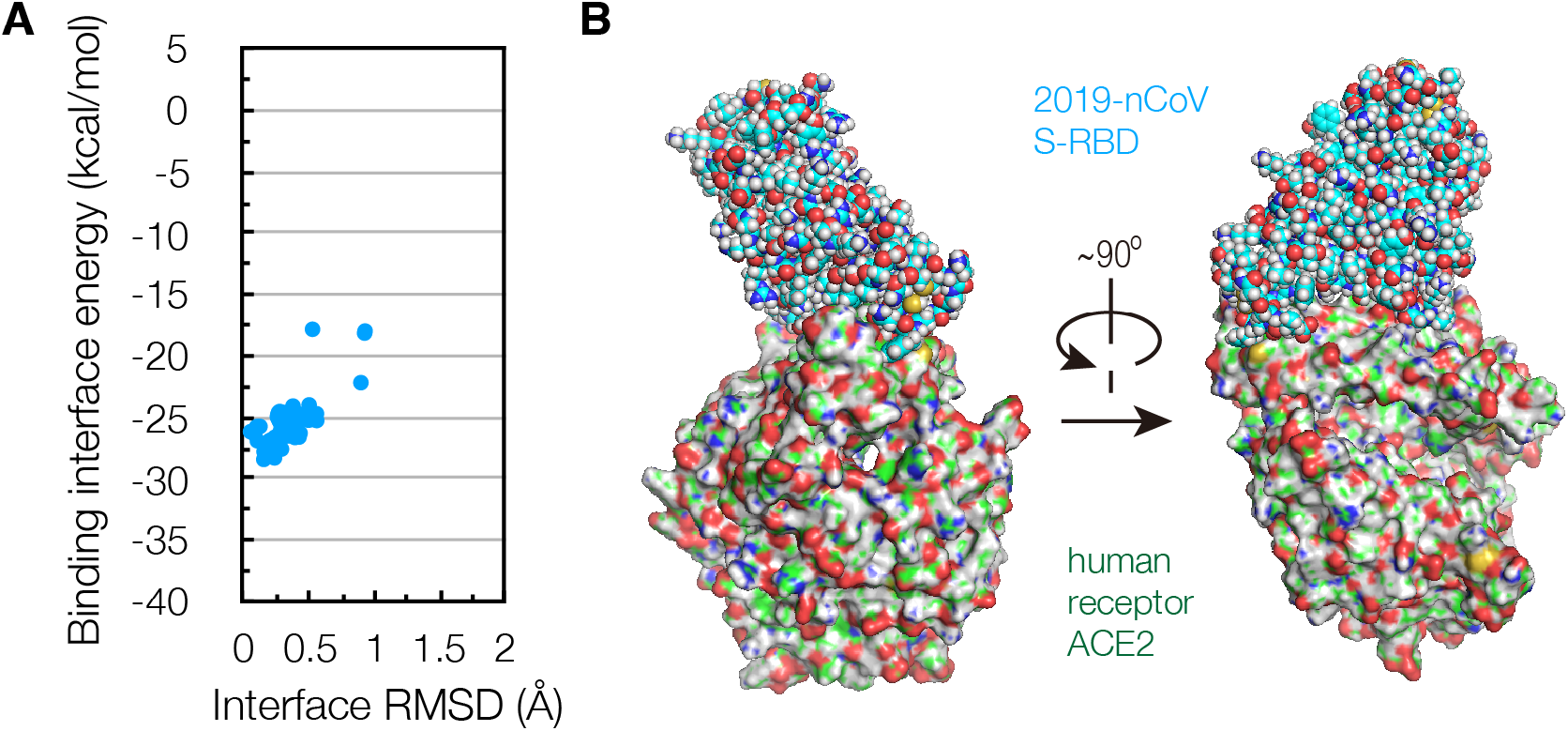
All-atom refinements of the lowest-energy binding model. (A) Scatter plot of binding free-energy scores versus RMSDs of interface residues of the refined models, where the RMSD reference is the lowest-energy conformation in Fig. 3A. (B) The lowest-energy model of S-RBD binding to ACE2 with an energy score of −28.4 kcal·mol^-1^.

### Comparison of human receptor-binding capability of 2019-nCoV with SARS-CoV

The successful identifications of the native-like binding conformations of S-RBDs to ACE2 enable a comparison of the receptor-binding capability of 2019-nCoV with that of SARS-CoV. Especially, the obtained energy scores based on the experimentally determined structure of SARS-CoV defined a binding free-energy scale, on which the relative receptor-binding strength of a newly emerged CoV could be evaluated.

As mentioned, according to statistical thermodynamics, the sampled binding conformations obtained in the Monte Carlo docking simulations can be considered to be thermodynamically probable in the binding process of S-protein to ACE2. Indeed, the binding free-energy (or binding affinity) is defined as a thermodynamic average of the binding energies over all possible binding modes of two interacting components. Therefore, the binding free-energy is approximated by the thermodynamic average of the binding energies of all the sampled low-energy conformations as:

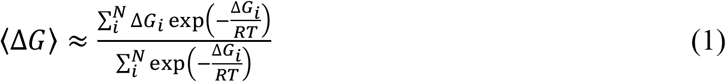

where *N* is the total number of the low-energy conformations sampled in the docking simulations, *R* is the ideal gas constant, and *T* is the absolute temperature (i.e., 298.15 K for the room temperature). Note that, when *N* approaches the number of all thermodynamically probable conformations, 〈Δ*G*〉 becomes the exact value.

With equation 1 and all the sampled binding conformations for SARS-CoV and 2019-nCoV, the binding free-energy of SARS-CoV S-RBD to ACE2 is:

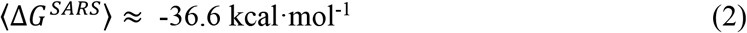

and that of 2019-nCoV S-RBD to ACE2 is:

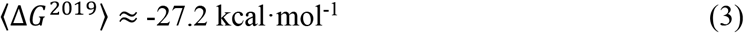

By comparing these two values, we found that 2019-nCoV S-RBD reaches about 73% of the binding strength of SARS-CoV S-RBD to ACE2. Since the receptor-binding of S-protein is the major driving force for the viral transmission to the human host cells, such a strength order supports that 2019-nCoV is able to infect the human host cells like SARS-CoV(Zhou et al., 2020).

Soon after the 2019-nCoV genome sequencing, Xu et al. (2020) also used structural modeling approach to predict the binding affinity of S-RBD to ACE2 In their study, two single complex conformations (i.e., one crystal structure and one predicted structure) were employed to calculate the binding free-energies of two CoVs, and showed that the affinity of 2019-nCoV S-RBD to ACE2 was in the order of ~65% of that for SARS-CoV. So, their predicted strength is about 10% lower than our value. Certainly, although our binding free-energy scores were calculated using a large-number of thermodynamically probable, low-energy binding conformations, further experimental measurements will be needed to testify the accuracy of both methods.

## Conclusions

We have performed large-scale protein-protein docking to evaluate the binding affinities of SARS-CoV and 2019-nCoV S-RBDs to the human receptor ACE2, in order to quantitatively assess the human receptor-binding capability of 2019-nCoV. Using the well-investigated SARS-CoV as the reference, our results showed that 2019-nCoV reaches about 73% of the ACE2 receptor-binding strength of SARS-CoV. This supports that the 2019-nCoV S-protein encoded by its genome has processed human receptor-binding ability close to that of SARS-CoV, and thus this newly emerged virus is likely able to bind to ACE2 and then to drive human-to-human transmission. However, the docking sampling for a large number of thermodynamically probable conformations did not find any binding conformation with an energy score lower than that of the lowest-energy conformation of SARS-CoV derived from experimentally determined structure, offering molecular evidence that the receptor-binding ability of 2019-nCoV is less than that of SARS-CoV.

Of course, ultimate confirmation of computational predictions requires experimental verification, which will also be very important for improving our method in the future. However, experimental approaches for deeply understanding the human transmission of viruses in terms of receptor-binding interactions and specificity is rather time consuming. Sometimes, it might be even impossible to employ experimental methods to thoroughly examine every evolved, mutant strain of a newly emerged virus, such as 2019-nCoV in our case. As an alternative, using a high-performance computer, our approach may assess the receptor-binding capability of a newly identified strain within a very short period of time. Thus, our method can be useful for the post-genome analysis for a newly identified CoV and its mutant strains, with the first-determined genome data. Note that, although viral genome data provide useful sequence information for comparison-based assessments with previously identified viruses, these data themselves do not offer hints about how the encoded viral proteins carry out their functions which may lead to serious human infections and diseases. Hence, post-genome analysis of viral protein functions is valuable not only for understanding the viruses themselves, but also for offering useful scientific supports for making effective, early decisions about viral prevention and control.

In summary, this study presents a fast method for accurately assessing the human transmission capability of newly emerged CoVs, and demonstrates that post-genome analysis of CoV protein-protein interactions may provide early scientific guidance for the viral prevention and control.

## Methods

### Homology modeling of 2019-nCoV S-RBD structure

The amino-acid sequence of 2019-nCoV S-protein was taken from the first genome sequence of a 2019-nCoV isolate (Wuhan-Hu-1) deposited in GenBank on January 12, 2020 (GenBank No: MN908947.3)(Wu et al., 2020). The initial structure of S-RBD was built according to standard modeling procedure on the SWISS-MODEL server at https://swissmodel.expasy.org, by submitting its amino-acid sequence in Fig. 2A and then selecting the SARS-CoV S-RBD structure (PDB 2AJF) as the modeling template. With the obtained model, high-resolution all-atom refinements were then carried out to optimize the conformations of the protein backbone and side-chains with the `relax` program of the Rosetta software suite(Das and Baker, 2008). And the Rosetta all-atom energy function 2015 (REF15)(Alford et al., 2017) was employed to calculate the conformational energy. The all-atom relax protocols are:

~~~
relax.linxgccrelease /
 –database $database_path /
 –s $start_model.pdb /
 –relax:thorough /
 –nstruct $model_number
~~~

### Local protein-protein docking

The `docking` program of Rosetta(Das and Baker, 2008) was used to carry out the local protein-protein docking with the starting conformation of a given S-RBD-ACE2 complex. The Rosetta energy function 2015 (REF15) was employed in the docking to evaluate the interaction energy of two docking components (i.e., S-RBD and ACE2). For each S-RBD-ACE2 pair, at least 3,000 independent docking simulations were performed to generate a large number of thermodynamically probable, low-energy binding conformations for calculating the binding free-energy. The top-scoring, lowest-energy conformations were identified as the native-like conformations of the S-RBD binding to ACE2. The local protein-protein docking protocols are:

~~~
docking_protocol.linxgccrelease /
 –database $database_path /
 –s $start_complex.pdb /
 –partners $ACE_chainID_$S-RBD_chainID /
 –dock_pert 3 8 /
 –ex1 /
 –ex2aro /
 –nstruct $model_number
~~~

### All-atom refinement of the lowest-energy conformation

The `docking` program of Rosetta was used to carry out the high-resolution refinement for further optimizing the lowest-energy binding conformation, with the Rosetta energy function 2015 (REF15). At least 200 independent docking runs were performed to generate the refined models. The all-atom refinement protocols are:

~~~
docking_protocol.linxgccrelease /
 –database $database_path /
 –s $lowest_complex.pdb /
 –docking_local_refine /
 –use_input_sc /
 –ex1 /
 –ex2aro /
 –nstruct $model_number
~~~

## Acknowledgements

The methodology development in this work was partly supported by the grant from the National Natural Science Foundation of China (No. 91430112).

## Additional information

Correspondence and requests for materials should be addressed to Q.H. or A.H.

